# Revisiting the reduction of stochastic models of genetic feedback loops with fast promoter switching

**DOI:** 10.1101/657718

**Authors:** J. Holehouse, R. Grima

**Affiliations:** University of Edinburgh

## Abstract

Propensity functions of the Hill-type are commonly used to model transcriptional regulation in stochastic models of gene expression. This leads to an effective reduced master equation for the mRNA and protein dynamics only. Based on deterministic considerations, it is often stated or tacitly assumed that such models are valid in the limit of rapid promoter switching. Here, starting from the chemical master equation describing promoter-protein interactions, mRNA transcription, protein translation and decay, we prove that in the limit of fast promoter switching, the distribution of protein numbers is different than that given by standard stochastic models with Hill-type propensities. We show the differences are pronounced whenever the protein-DNA binding rate is much larger than the unbinding rate, a special case of fast promoter switching. Furthermore we show using both theory and simulations that use of the standard stochastic models leads to drastically incorrect predictions for the switching properties of positive feedback loops and that these differences decrease with increasing mean protein burst size. Our results confirm that commonly used stochastic models of gene regulatory networks are only accurate in a subset of the parameter space consistent with rapid promoter switching.

**Statement of Significance:** A large number of models of gene regulatory networks in the literature assume that since promoter switching is fast then transcriptional regulation can be effectively modeled using Hill functions. While this approach can be rigorously justified for deterministic models, it is presently unclear if it is also the case for stochastic models. In this article we prove that this is not the case, i.e. stochastic models of gene regulatory systems, namely those with feedback loops, describing transcriptional regulation using Hill functions are only valid in a subset of parameter conditions consistent with fast promoter switching. We identify parameter regimes where these models are correct and where their predictions cannot be trusted.

## 1 Introduction

Many biochemical systems have one or more species with low molecule numbers which implies that the dynamics can be highly noisy and consequently a deterministic description may not be accurate [1–4]. Rather a more appropriate mathematical description is stochastic and given by the chemical master equation [5]. When the system is made up of zero and first order reactions only, exact solutions at both steady state and in time are occasionally possible [6]. However many systems have at least one bimolecular reaction and in such cases only a few exact steady-state solutions of the CME are known (see for example [7–10]). A common example of such systems are auto-regulatory feedback loops, whereby a protein produced by a gene binds to its own promoter region to activate or suppress its own production [11–13]. In the absence of exact solutions, we become either reliant: (i) on the stochastic simulation algorithm (SSA) [14] or (ii) on approximations of the original network so that analytic results become tractable [15–17]. Generally, it is a challenge to utilise approximations to simplify the CME such that the resulting reduced equation is representative of the true system dynamics.

A common set of approximation methods are based on timescale separation. At the microscopic level, there are several different scenarios which can lead to timescale separation conditions. Depending on their firing rate, reactions can be classified as either slow or fast. Depending on the reaction system, it is possible that fast and slow reactions do not involve the same species but more commonly fast and slow reactions share some species and hence it is generally unclear what should be considered a fast or a slow species. Methods in the literature differ according to the definition of what is a slow or fast species. Zeron and Santillan [18] assume that fast species are only involved in fast reactions whereas slow species can participate in both slow and fast reactions. In contrast, Cao et al. [19] define slow species as those involved in slow reactions only and fast species as those participating in at least one fast reaction and any number of slow reactions. These two approaches lead to a reduced CME description in the slow species only. Other approaches due to Haseltine and Rawlings [20] and Goutsias [21] model the state of the system using extents of reaction as opposed to molecules of species. A singular perturbation theory based method has also recently been used to obtain a reduced stochastic description [22]. There are also several formal results that have been mathematically proven for reaction systems in various scenarios [23–26].

Despite the wide breadth of rigorous approaches (e.g. perturbation theory [27]), by far the most popular approach in the literature of computational and systems biology to obtain a reduced master equation is heuristic. The key idea is to use the results of timescale separation for deterministic kinetics. Under the quasi-steady state or fast equilibrium approximations, the mean concentration of a subset of species (the fast species) reaches steady-state on a much shorter time scale than the rest of the species (the slow species). Using the deterministic rate equations it is then possible to express the concentration of the fast species in terms of the concentration of the slow species. This leads to a reduced chemical system composed of effective reactions with non-mass action kinetics describing the dynamics of the slow species. The reduced chemical master equation is then obtained by writing effective propensities analogous to the non-mass action reaction rates obtained from the deterministic analysis. For example, Hill-type effective protein production rates in the deterministic rate equations result if the gene equilibrates on a much faster timescale than mRNA and protein, i.e the fast promoter switching limit (see for example [28] for experimental evidence of this limit), and hence by analogy, Hill-type propensities for the protein production rates are commonly used in stochastic simulations of gene regulatory networks [29–36]. All of these studies and many others assume that such effective propensities are justified in the limit of fast promoter switching.

The advantage of this heuristic approach is its simplicity and ease of use and this is the main reason for its widespread use. However clearly the use of a reduced master equation obtained from deterministic considerations is doubtful. This has led to a number of studies evaluating the accuracy of these reduced master equations. Thomas et al. in a series of papers [37–39] showed using Langevin approximation theory that in the limit of large molecule numbers and for parameters consistent with the quasi-steady state approximation, the mean number of molecules of slow species predicted by the reduced master equation agrees with that predicted from the master equation of the full system but the variance of molecule number fluctuations does not. Similar results have been shown using stochastic simulations by Kim et al. (in particular see Fig. 1 of [40]). In contrast, Bundschuh et al. [41] have shown that the SSA corresponding to the heuristic reduced master equation of a negative feedback loop, whereby the DNA-protein binding reactions are assumed fast compared to the rest of the reactions and hence are eliminated, is in very good agreement with the SSA of the full system for parameter values specific to the phage *λ* system (this case is referred to as “Michaelis-Menten system” in their paper). At first sight the results of Thomas et al. and Bundschuh et al. may appear contradictory but in reality they are not: while the results of Thomas et al. prove that the heuristic approach of obtaining reduced master equations cannot be considered equivalent to the stochastic version of the quasi-steady state approximation (see also [42]), nevertheless it is possible that the error in the predictions of the heuristic approach are small for specific parameter values which would be consistent with the results of Bundschuh et al. What is clear from these studies is that more work is needed to identify the precise regions of parameter space where the heuristic reduced master equation of gene regulatory networks can be safely used – the study reported in this article identifies such regions and hence fills a gap in the literature.

**Figure 1:**
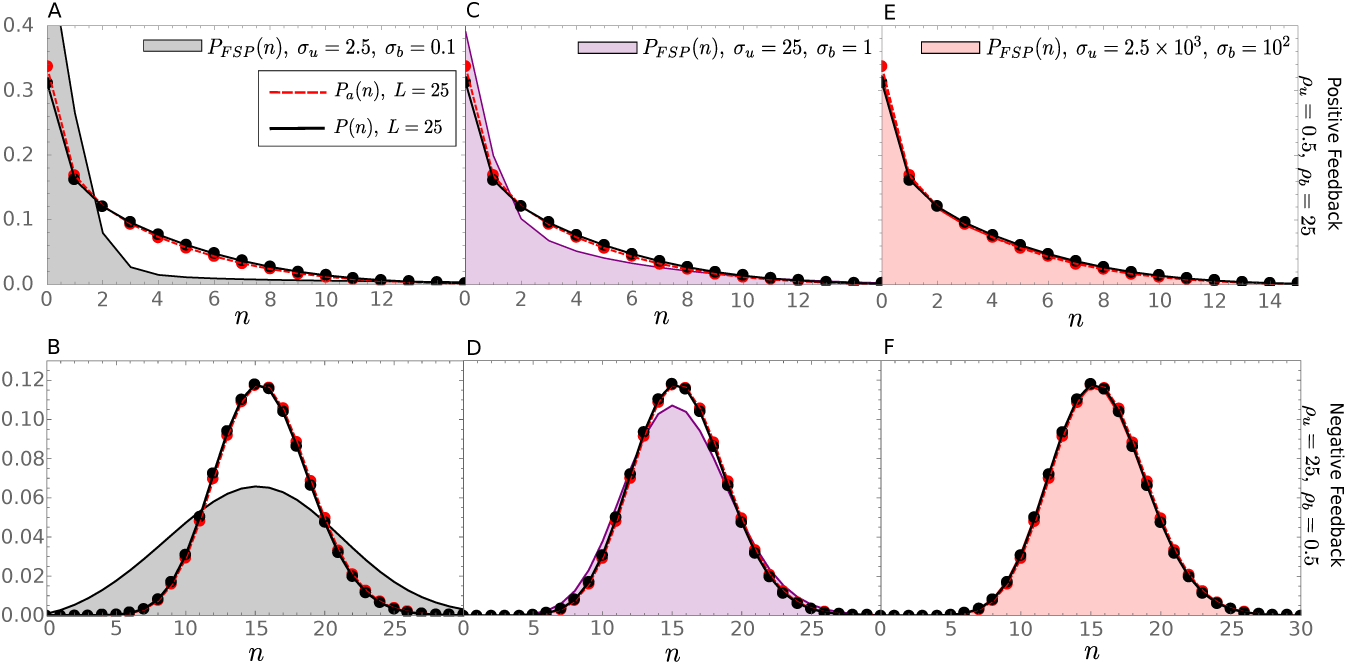
Plots comparing the probability distributions of proteins as predicted by the heuristic reduced master equation (*P*_*a*_ (*n*)), the exact reduced master equation (*P*(*n*)) and the full master equation in the limit of large *L* (*P*_*FSP*_ (*n*)). *L* is kept constant throughout the figure. Shaded regions indicate the solution of the full master equation Eq. (27) using FSP, dashed red lines indicate the heuristic probability distribution from Eq. (11), and black solid lines indicate the exact solution in the fast promoter switching limit from Eq. (28). Throughout the paper FSP is used as the benchmark for our analytic results, with a state space truncation chosen such that the probability distributions are indistinguishable from SSA. Going from left-to-right one can observe how Eqs. (11) and (28) correctly describe the large *L* limit (here *L* = 25) when both *σ*_*b*_ and *σ*_*u*_ are themselves large, i.e. the fast promoter switching limit. The top row of plots show this for the case of positive feedback (*ρ*_*u*_ = 0.5 and *ρ*_*b*_ = 25) and the bottom row of plots show this for negative feedback (*ρ*_*u*_ = 25 and *ρ*_*b*_ = 0.5).

The structure of this article is as follows. In Section 2 we obtain the steady-state solution of the heuristic reduced master equation with Hill-type protein production propensities for a non-bursty genetic feedback loop and prove that it is different than the solution of the master equation of the non-bursty genetic feedback loop in the limit of fast promoter switching. It is then shown that the differences between the probability distributions of protein numbers predicted by the two master equations tend to zero when the rate of DNA-protein binding is much smaller than the unbinding rate. In contrast, the differences maximize when the rate of DNA-protein binding is much larger than the unbinding rate. The results are confirmed by stochastic simulations across large swathes of parameter space. In Section 3 we extend the analysis to bursty feedback loops. We finish with Section 4, providing a discussion and then by concluding our results.

## 2 Model reduction for non-bursty feedback loops

### 2.1 Deterministic description and reduction

The reaction scheme for a genetic non-bursty feedback loop is given by:

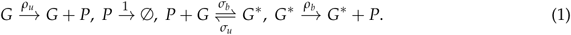

This models the production of proteins, their degradation, DNA-protein binding and unbinding. For simplicity we do not have an mRNA description (though this will be added in Section 3). The gene can be in one of two states: an unbound state *G* and a bound state *G*\*. The rate of protein production depends on the gene state. Note that here we have scaled all parameters by the protein degradation rate. Of course, given that there is only one copy of the gene, one expects fluctuations to be important and a stochastic model to be the most appropriate mathematical description. However for the moment we shall ignore the inherent stochasticity and analyze the system using a deterministic approach. The deterministic rate equations are:

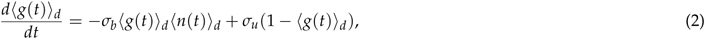

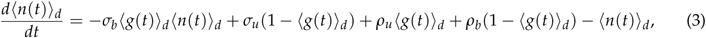

where ⟨*n*(*t*)⟩_*d*_ denotes the mean number of molecules of protein *P* at time *t*, ⟨*g*(*t*)⟩_*d*_ denotes the mean number of molecules of gene *G* at time *t* and *g*\*⟨(*t*) ⟩_*d*_ denotes the mean number of molecules of gene *G*\* at time *t*. Since the gene can only be in either the bound or unbound state at any one time, one may also interpret ⟨*g*(*t*)⟩_*d*_ and ⟨*g*\*(*t*)⟩_*d*_ as the mean fraction of time spent in either gene state respectively. These mean molecule numbers are calculated within the deterministic approximation (hence the subscriptd) and will generally be different than the mean molecule numbers of the system obtained from a stochastic description of the system [43]. Note that we have used the relation ⟨*g*(*t*)⟩_*d*_ + ⟨*g*\* (*t*)⟩_*d*_ = 1, i.e. there is one gene copy. Note also that *t* is non-dimensional time, i.e. actual time multiplied by the protein degradation rate. It can also be shown (see Appendix A) that the deterministic equations Eqs. (2)–(3) agree with those from the chemical master equation under the assumption of independence of fluctuations in the protein and gene numbers.

By the fast equilibrium approximation it follows that *∂*_*t*_⟨*g*(*t*)⟩_*d*_ ≈ 0 (and *∂*_*t*_⟨*g*\* (*t*)⟩_*d*_ ≈ 0) for all times which implies, from Eq. (2), that ⟨*g*(*t*)⟩_*d*_ = *L*/(*L* + ⟨*n*(*t*) ⟩_*d*_) where *L* = *σ*_*u*_ /*σ*_*b*_. The definition of *L* is used frequently throughout the text. Substituting the latter in the right hand side of Eq. (3) and suppressing the time dependence (for notational convenience) we obtain:

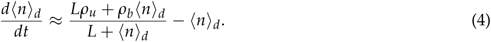

This is an effective time-evolution equation for the protein numbers, within the deterministic approximation. This corresponds to a system with two reactions: an effective zero-order reaction modeling the transcriptional regulation of protein production, and a first-order protein degradation reaction. The rate of protein production is a function of the mean number of proteins and three special cases can be distinguished: (i) If *ρ*_*u*_ > *ρ*_*b*_ then the rate of protein production decreases with increasing ⟨*n*⟩_*d*_; this is the case of negative feedback. (ii) If *ρ*_*u*_ < *ρ*_*b*_ then the rate of protein production increases with in-creasing ⟨*n*⟩_*d*_; this is the case of positive feedback. (iii) If *ρ*_*u*_ = *ρ*_*b*_ then the rate of protein production is independent of ⟨*n*⟩_*d*_ and effectively there is no feedback.

Intuitively one would expect the solution of Eq. (4) to be an excellent approximation to the time-evolution of the protein in the full model given by Eqs. 2–3), in the limit of fast promoter switching, i.e. min(*σ*_*u*_, *σ*_*b*_) ≫ max(1, *ρ*_*u*_, *ρ*_*b*_). This can be explicitly verified by calculating the ratio of gene and protein timescales, as follows. Replacing *σ*_*u*_ by *σ*_*u*_ /*ϵ* and *σ*_*b*_ by *σ*_*b*_ /*ϵ* and taking the limit of *ϵ* → 0, it is straightforward to show that to leading order the two eigenvalues of the Jacobian matrix of the rate equations Eqs. 2–3) evaluated at steady-state, are given by:

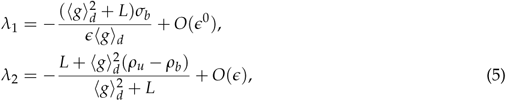

where ⟨*g*⟩_*d*_ is the steady-state mean gene number given by:

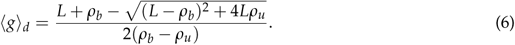

For completeness and since we will use it later, the steady-state mean protein number is given by:

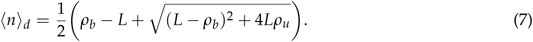

Note that *λ*_1,2_ are negative and hence the steady-state of the system is stable to small perturbations. Furthermore Eq. (5) shows that as *ϵ* → 0, *λ*_1_ → −∞ and *λ*_2_ tends to a constant. Since the timescales of decay of transients in the mean protein and gene numbers are given by the absolute of the inverse of the eigenvalues, it follows that there is clear timescale separation in the limit of fast promoter switching. Note that in the calculation above we assumed that *ρ*_*u*_ ≠ *ρ*_*b*_; a similar calculation for the equality case also leads to timescale separation.

Hence to summarize, a deterministic rate equation analysis shows that in the limit of fast promoter switching, the reaction scheme (1) composed of five reactions (four first-order reactions and a bimolecular reaction) reduces to just two reactions: an effective zero-order reaction for the production of proteins with a rate which is a function of the mean number of proteins and a first-order reaction modeling protein degradation.

### 2.2 Heuristic stochastic model reduction

As mentioned in the Introduction, one of the most popular stochastic model reduction approaches consists of directly writing the chemical master equation for the reduced reaction scheme deduced from the deterministic analysis in Section 2.1. In particular given there are *n* proteins in the system then we define the effective propensities:

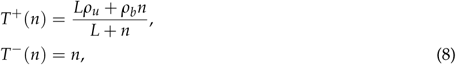

where *T*^+^(*n*)*dt* is the probability, given *n* proteins, that a protein production reaction increasing the number of proteins by one, will occur in the time interval [*t, t* + *dt*) and *T*^−^ (*n*)*dt* is the probability, given *n* proteins, that a protein degradation event reducing the number of proteins by one will occur in the time interval [*t, t* + *dt*). These probabilities are deduced directly from the form of the effective rate equation Eq. (4). Essentially the probability per unit time for a particular reaction is taken to be the same as the reaction rate in the effective deterministic rate equation with ⟨*n*⟩ replaced by *n*. The chemical master equation for this reduced reaction scheme is then given by:

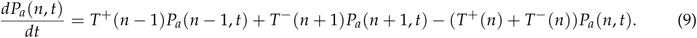

Note that we have labelled the solution of this *approximate* heuristic master equation *P*_*a*_ to distinguish it from the solution of the full master equation *P*, which we discuss in Section 2.4. The equations for the mean number of protein ⟨*n*(*t*)⟩_*a*_ = Σ_*n*_ *nP*_*a*_ (*n, t*) can be derived from the master equation:

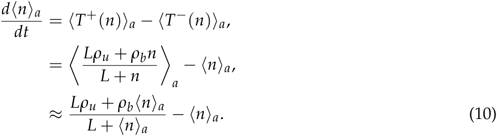

Note that in the last line we have made use of the fact that in the limit of small protein number fluctuations *n* can be replaced by its average. *Hence while the selection of the propensities stems from a heuristic rule with no fundamental microscopic basis, nevertheless it guarantees equivalence between the effective equation for the time-evolution of the mean protein numbers of the heuristic master equation and the reduced deterministic rate equation in the limit of small protein number fluctuations (since Eq. (4) and Eq. (10) are the same upon interchanging* ⟨*n*⟩_*d*_ *by* ⟨*n*⟩_*a*_). Note that however for the general case of non-vanishing protein fluctuations, the mean of the heuristic stochastic model is different than that predicted by the deterministic rate equations.

The exact solution of the one variable master equation Eq. (9) in steady-state conditions can be obtained using standard methods [44] and is given by:

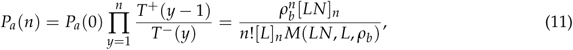

where *N* = *ρ*_*u*_ /*ρ*_*b*_, [*x*]_*n*_ = *x*(*x* + 1)…(*x* + *n* − 1) (the Pochhammer symbol) and *M* is the Kummer confluent hypergeometric function. The definition of *N* is used frequently throughout the text. To obtain insight into the discrepancies introduced by the heuristic approach, we now study two limiting cases.

#### 2.2.1 The limit of large *L*

This is the limit in which the rate at which proteins bind DNA is much smaller than the unbinding rate. In this limit, the propensities given by Eq. (8) reduce to the simpler form:

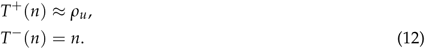

Hence in this limit, the propensity *T*^+^(*n*) is independent of *n* and the steady-state solution of Eq. (9) is simply a Poisson with mean *ρ*_*u*_:

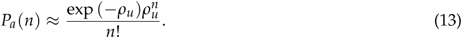

Note that this derivation is intuitive but not formally precise because we have implicitly assumed the exchange of two limits: lim_*L*_→_∞_ lim_*t*→∞_ *P*_*a*_ (*n, t*) = lim_*t*→∞_ lim_*L*_→_∞_ *P*_*a*_ (*n, t*). A formal proof of this result starting from the exact solution Eq. (11) can be found in Appendix C.

#### 2.2.2 The limit of small *L*

This is the limit in which the rate at which proteins bind DNA is much larger than the unbinding rate. In this limit, the propensities given by Eq. (8) reduce to the simpler form:

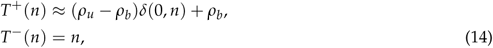

where *δ*(0, *n*) is the Kronecker delta. Substituting these in the heuristic reduced master equation Eq. (9), multiplying throughout by *z*^*n*^ and taking the sum over *n* on both sides of this equation we get the corresponding generating function equation:

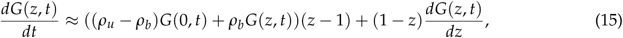

where *G*(*z*) = Σ_*n*_ *z*^*n*^ *P*_*a*_ (*n, t*). In steady-state, this equation has the solution:

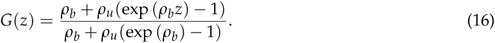

Hence the steady-state probability distribution is given by:

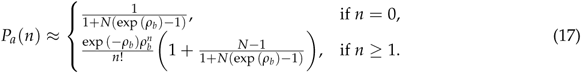

While intuitive, this proof suffers from the same looseness with exchange of limits as in Section 2.2.1. An alternative rigorous proof of this result starting from the exact solution Eq. (11) can be found in Appendix C. Eq. (17) is clearly not a Poisson when there is positive or negative feedback (*N* ≠ 1). Note that when *N* = 1, the solution is a Poisson but this case is biologically unimportant because it implies that the rate of protein production is independent of the state of the promoter (bound or free) and consequently there is effectively no feedback mechanism at play.

### 2.3 Conditions for the validity of heuristic stochastic model reduction

To obtain further insight into the conditions under which the heuristic model reduction is correct, we consider the stochastic description of the non-bursty feedback loop (1) but ignoring fluctuations in the protein numbers stemming from the reversible binding of protein to gene. The neglection of protein binding fluctuations corresponds to the following reaction scheme:

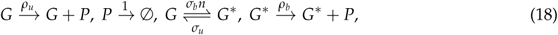

where *n* in the reaction rate denotes the number of free proteins. This is a common approximation in the literature [36, 45, 46], the rationale being that since protein numbers are typically much larger than one hence the gain or loss of one molecule via the gene binding reactions can be safely ignored. Given this assumption, the chemical master equation of the non-bursty feedback loop (1) can be conveniently written as a set of two coupled equations:

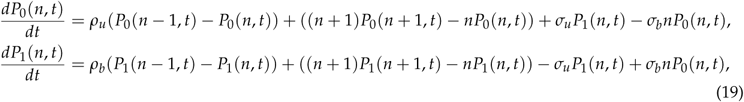

where *P*_0_(*n, t*) is the probability that at time *t* there are *n* proteins and the gene is in state *G* while *P*_1_(*n, t*) is the probability that at time *t* there are *n* proteins and the gene is in state *G*\*. Note that time *t* is non-dimensional and equal to the actual time multiplied by the protein degradation rate. The probability of *n* proteins is then given by *P*(*n, t*) = *P*_0_(*n, t*) + *P*_1_(*n, t*). Defining the generating functions *G*_0_(*z, t*) = Σ_*n*_ *z*^*n*^ *P*0(*n, t*) and *G*_1_(*z, t*) = Σ_*n*_ *z*^*n*^ *P*_1_(*n, t*), the generating function differential equations corresponding to Eqs. (19) are given by:

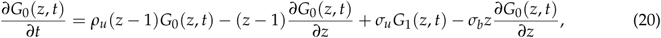

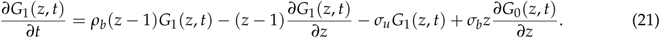

We can solve for the total generating function *G*(*z*) = *G*_0_(*z*) + *G*_1_(*z*) as follows. At steady state *∂G*_*i*_ (*z, t*)/*∂t* = 0, and utilising the relation *G*_1_(*z*) = *G*(*z*) − *G*_0_(*z*) we can use the sum of Eq. (20) and (21) to find *G*_0_(*z*) = *G*_0_(*G*(*z*), *G*′(*z*), *z*) and 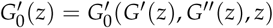 below

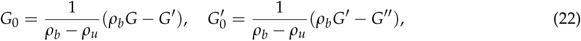

where we suppress the *z* dependence for brevity. We then substitute Eq. (22) into Eq. (20), again using *G*_1_ = *G* − *G*_0_, to give a second order linear differential equation in terms of *G*,

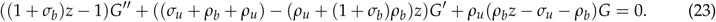

This differential equation has two singularities, a regular singularity at *z* = 1/(1 + *σ*_*b*_) and an irregular singularity at *z* = ∞ and hence its solution is amenable to the _1_*F*_1_(*α*; *β*; *z*) hypergeometric function (otherwise known as the Kummer function *M*(*α*; *β*; *z*)). Using a change of variable and an exponential transformation we confirm this, and the solution is given as

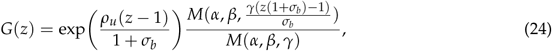

Where

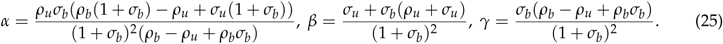

In the limit of fast promoter switching, i.e. replacing *σ*_*u*_ by *σ*_*u*_ /*ϵ* and *σ*_*b*_ by *σ*_*b*_ /*ϵ* and taking the limit of *ϵ* → 0, one can show that the leading-order term in the series expansion of Eq. (24) in powers of *ϵ* is given by:

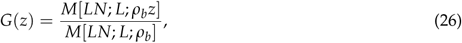

where we remind the reader of definitions *L* = *σ*_*u*_ /*σ*_*b*_ and *N* = *ρ*_*u*_ /*ρ*_*u*_. It is easy to show that *P*(*n*) = (1/*n*!)*d*^*n*^ *G*(*z*)/*dz*^*n*^|_*n*=0_ precisely equals Eq. (11).

Hence we have shown that if protein number fluctuations due to reversible binding can be ignored then the stochastic description agrees with that of the heuristic master equation in the limit of fast promoter switching. This indeed gives some credibility to the use of the heuristic master equation and explains the widespread belief, based on stochastic simulations, that the heuristic master equation is correct in the limit of fast promoter switching. This result is however surprising when one considers that the heuristic is often justified from deterministic arguments, and that the deterministic treatment ignores the sizable fluctuations associated with gene switching.

### 2.4 Exact stochastic model reduction

The master equation we solved in the previous section is not the exact master equation since we have ignored protein binding fluctuations. In what follows we properly take these into account. For the non-bursty feedback loop (1), the stochastic description is given by the chemical master equation which can be conveniently formulated as a set of two coupled equations:

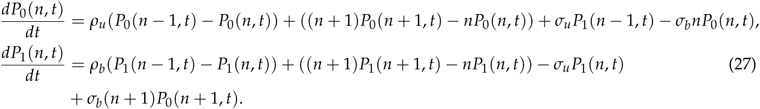

Note that these equations are the same as Eq. (19) except for the terms describing protein-gene binding, i.e. those proportional to *σ*_*b*_ and *σ*_*u*_. In the limit of fast promoter switching, i.e., replacing *σ*_*u*_ by *σ*_*u*_ /*ϵ* and *σ*_*b*_ by *σ*_*b*_ /*ϵ* and taking the limit of *ϵ* → 0, one can show that the steady-state solution of Eqs. (27) (to leading-order in *ϵ*) is given by:

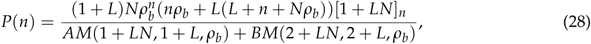

where

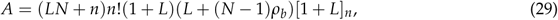

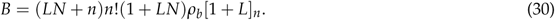

See Appendix B for the details of the derivation. Comparing Eq. (28) with the solution of the heuristic reduced master equation Eq. (11), it is immediately obvious that the two are not equal. *Hence it follows the heuristic reduced master equation is generally different than the solution of the full master equation under fast promoter switching conditions, contradicting a major assumption in the literature (as discussed in the Introduction)*. It is straightforward to verify that the two agree only if *N* = 1 in which case the protein production rate is the same in state *G* or *G*\* and hence there is no effective feedback mechanism.

To understand the nature of the differences between Eq. (11) and Eq. (28), we consider two limiting cases of small and large *L*. Whilst results in these limits can be obtained directly from consideration of Eq. (28) (see Appendix C) it is both simpler and instructive to consider a different approach which does not need the exact solution of the master equation. The advantage of this approach is that as we shall see later on, it can be easily extended to the analysis of more complex feedback systems.

Since ⟨*g*⟩ is the fraction of time spent in state *G* for the full stochastic model, it follows that in the limit of small *L*, ⟨*g*⟩ is also very small, the gene spends most of its time in state *G*\* and consequently the principal reactions determining the protein dynamics are 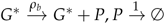. Similarly it can be argued that in the limit of large *L*, ⟨*g*⟩ ≈ 1 (the gene spends most of its time in state *G*) and hence the principal reactions determining the protein dynamics are 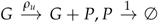. The master equation for both sets of principal reactions is trivial to solve and implies that the steady-state protein number distribution in both limits is a Poisson:

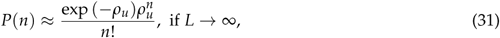

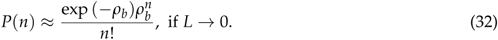

A formal derivation of these results starting from the solution Eq. (28) can be found in Appendix C.

### 2.5 Comparison of heuristic and exact reduction for small & large *L*

Comparing Eqs.(32)–(31) with Eq. (13) and Eq. (17), it is immediately clear that the heuristic method of stochastic model reduction gives the correct answer in the limit of fast promoter switching for large *L* but the incorrect answer for small *L*. Note that timescale separation exists in both cases of small and large *L* (as can be verified using Eq. (5)) and hence, the lack of agreement of the heuristic and exact reduction is not expected. In Fig. 1 we verify that the heuristic and exact reductions agree with each other and with the Finite State Projection (FSP) of the full master equation for large *L* provided the fast promoter switching limit (large *σ*_*u*_ and *σ*_*b*_ compared to all other parameters) is also met. This is the case for both positive and negative feedback. Note that FSP is a computationally efficient non Monte-Carlo method that solves the master equation to any desirable degree of accuracy [47].

To further understand the differences between the two protein distributions in the limit of small *L* we now look at the mean protein numbers, the Fano Factor (FF) and the Coefficient of Variation (CV) of protein number fluctuations:

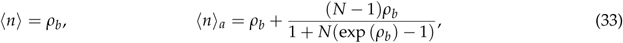

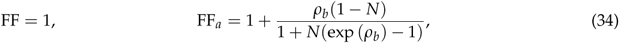

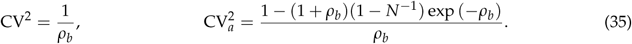

Note that the subscript *a* denotes calculation using Eq. (17) while no subscript implies calculation using Eq. (32).

From Eq. (33) we deduce that ⟨*n*⟩_*a*_ < ⟨*n*⟩ for *N* < 1 and ⟨*n*⟩_*a*_ > ⟨*n*⟩ for *N* > 1. This means that the solution of the approximate heuristic master equation *underestimates* the mean for positive feedback (*N* < 1) and *overestimates* the mean for negative feedback (*N* > 1). Since the deterministic rate equations also predict a steady-state protein mean of *ρ*_*b*_ for the case *L* → 0 (see Eq. (7)) it then follows that the approximate heuristic master equation also leads one to believe in noise-induced shifts of the mean which actually do not exist. From Eq. (34) we deduce that the approximate heuristic master equation artificially predicts *sub-Poissonian* (FF_a_ < 1) fluctuations in molecule numbers for negative feedback (*N* > 1) and *super-Poissonian* (FF_a_ > 1) fluctuations in molecule numbers for positive feedback (*N* < 1). These deviations from Poissonian behavior are most pronounced for intermediate *ρ*_*b*_ since for small and large *ρ*_*b*_, FF_*a*_ ≈ 1. From Eq. (35) we deduce that 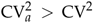 for *N* < 1 and 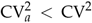 for *N* > 1, i.e. the approximate heuristic master equation overestimates the size of the protein number fluctuations for positive feedback and underestimates them for negative feedback. These observations are confirmed for positive feedback loops in Fig. 2. In particular note that in Fig. 2(A) the heuristic reduced master equation predicts switch-like behavior as *ρ*_*b*_ is increased (from zero for *ρ*_*b*_ below approximately 5 to larger than zero for *ρ*_*b*_ > 5) while the exact reduced master equation predicts no such transition for this set of parameters – the lack of reliability in predicting the switching characteristics of positive feedback loops is notable because previous studies [33] have used the heuristic reduced master equation to study switching phenomena.

**Figure 2:**
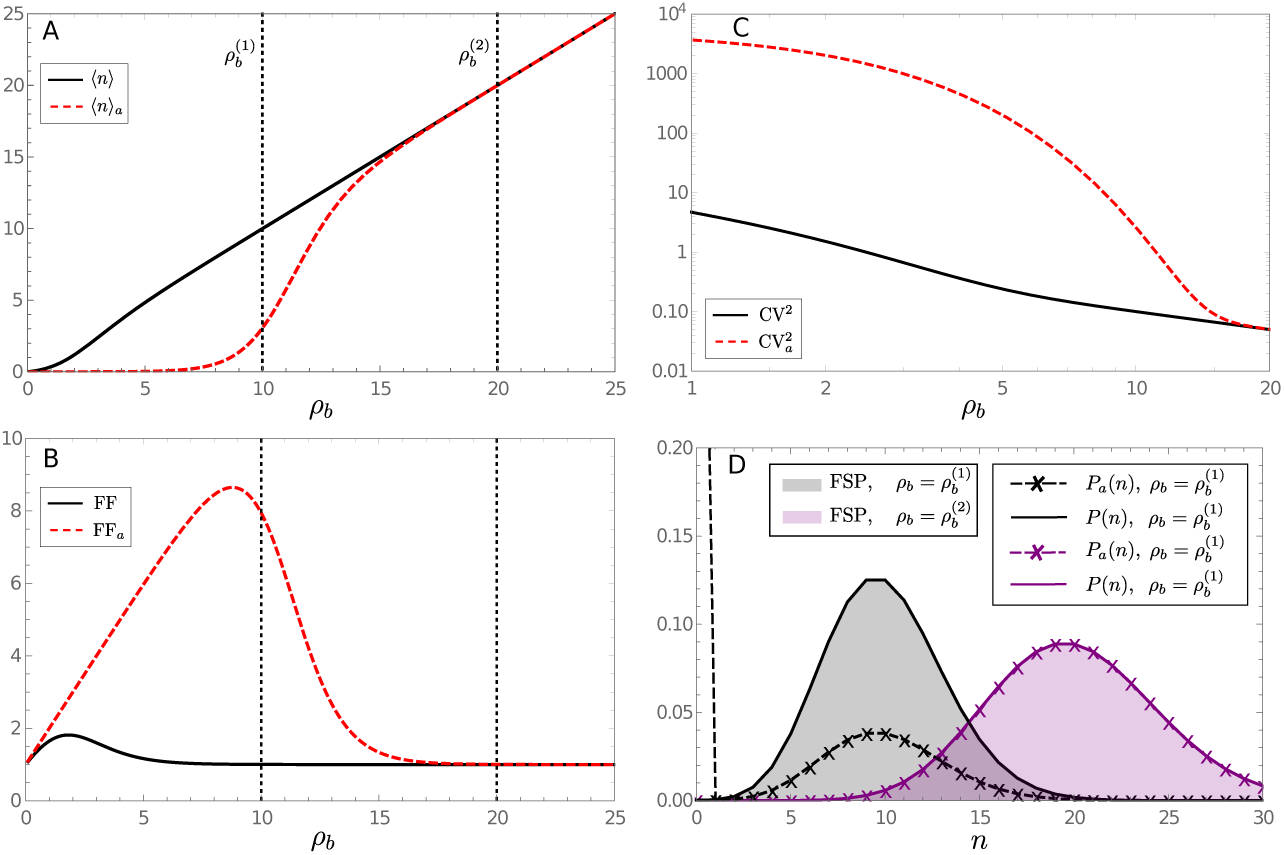
Plots showing the breakdown of the heuristic reduced master equation for fast promoter switching in the limit of small *L* and positive feedback in steady-state conditions. The plots show the mean protein number (A), the Fano Factor of protein number fluctuations (B) and the Coefficient of Variation squared (C) as a function of *ρ*_*b*_. In (D) we show the probability distribution of protein numbers corresponding to two different values of *ρ*_*b*_. The rest of the parameters are fixed to *σ*_*u*_ = 10^2^, *σ*_*b*_ = 10^5^ and *ρ*_*u*_ = 0.0002; this implies *L* = 10^−3^. Note that min(*σ*_*u*_, *σ*_*b*_) ≫ max(1, *ρ*_*u*_, *ρ*_*b*_) and hence fast promoter switching is ensured. Note that ⟨*n*⟩_*a*_, *FF*_*a*_ and 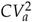 in (A)-(C) are calculated using the solution of the heurestic master equation Eq. (11) while their non-subscript versions are calculated using the solution of the exact reduced master equation Eq. (28). These are in good agreement with the moments calculated in the limit of small *L* and given by Eq. (33)–(35). The distributions *P*_*a*_ (*n*) and *P*(*n*) in (D) are calculated using Eq. (17) and Eq. (32), respectively. The plots verify the large differences between the heuristic and exact reduced master equation for positive feedback loops (see text for discussion), as well as showing agreement between the theoretical distribution for the exact reduced master equation and that obtained using FSP of the full master equation Eq. (67).

We can also show that generally positive feedback leads to larger deviations of the heuristic from the exact stochastic model reduction than is the case for negative feedback. Consider strong positive feedback *ρ*_*u*_ ≪ *ρ*_*b*_ (*N* ≪ 1) with the additional constraint *ρ*_*u*_ ≪ *ρ*_*b*_ exp (−*ρ*_*b*_). From Eq. (33) it can then be shown that ⟨*n*⟩ = *ρ*_*b*_, ⟨*n*⟩_*a*_ ≈ 0. If we now reverse the values of *ρ*_*u*_ and *ρ*_*b*_ such that we have strong negative feedback then *ρ*_*b*_ is very small and *N* ≫ 1 then ⟨*n*⟩ = *ρ*_*b*_, ⟨*n*⟩_*a*_ ≈ 1. Clearly |⟨*n*⟩ − ⟨*n*⟩_*a*_| is much larger for positive feedback than negative feedback (with *ρ*_*u*_, *ρ*_*b*_ interchanged) and this difference is evident in the distributions as well. Finally we compute the conditions for the existence of a mode of the probability distribution at *n* = 0 using Eq. (32) and Eq. (17) respectively:

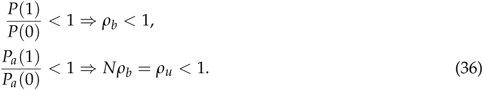

This implies that if *ρ*_*u*_ < 1, *ρ*_*b*_ > 1 (a special case of positive feedback), the approximate heuristic master equation predicts an artificial mode at *n* = 0 whereas if *ρ*_*u*_ > 1, *ρ*_*b*_ < 1 (a special case of negative feedback), the approximate heuristic master equation misses to predict an actual mode at *n* = 0. These predictions, contrasting the differences between the heuristic and exact model reduction for positive and negative feedback loops, are illustrated in Fig. 3. Note that multiple peaks in the protein distribution are often thought to describe switching between different phenotypes and hence of importance to understanding cellular decision-making [48, 49] – the lack of accuracy in the heuristic model predictions for the bimodality of the protein distribution shows that use of this model can lead to incorrect biological predictions.

**Figure 3:**
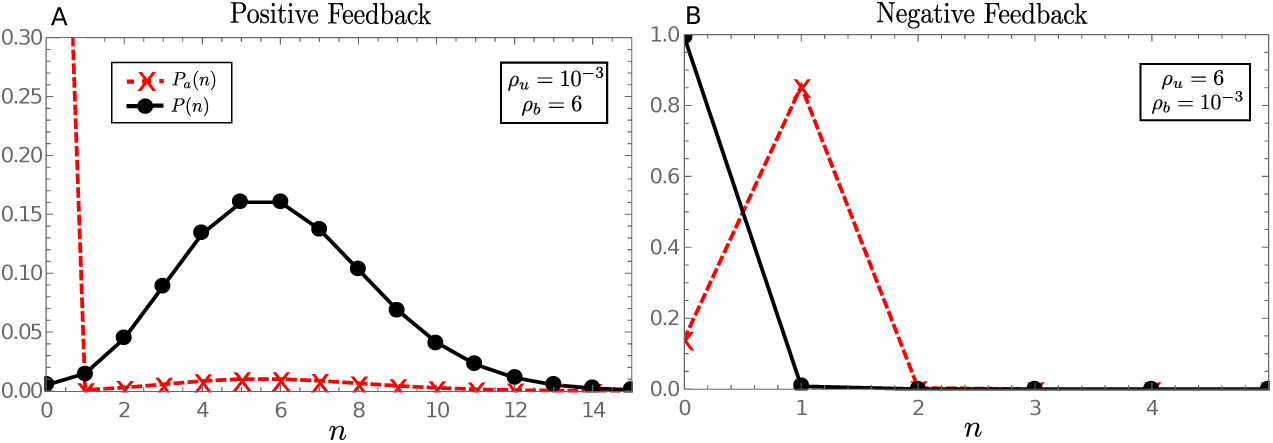
Plots comparing the steady-state protein distributions predicted by the heuristic and exact reduced master equations for strong positive feedback (A) and strong negative feedback (B) for the case of small *L. P*_*a*_ (*n*) is calculated using Eq. (11) while its subscript version is calculated using the solution of the exact reduced master equation Eq. (28). Note that the heuristic distribution (*P*_*a*_ (*n*)) predicts an artificial mode at zero for positive feedback and misses the prediction of a mode at zero for negative feedback, in line with the conditions Eq. (36). Note also that as predicted by theory, the differences between *P*(*n*) and *P*_*a*_ (*n*) are most significant for positive feedback loops since for negative feedback loops the differences amount to the order of a single molecule. The parameters *σ*_*u*_ = 10^2^ and *σ*_*b*_ = 10^5^ are fixed across both plots (implying *L* = 10^−3^) while the values of *ρ*_*u*_ and *ρ*_*b*_ are stated on the figure. FSP distributions are indistinguishable from the theoretical distributions shown in the figure.

The differences between the heuristic and exact model reduction can be explained using the results of Section 2.3. There we showed that the heuristic master equation has the same solution, in the limit of fast promoter switching, as the master equation which ignores protein fluctuations due to the reversible protein-DNA binding reaction.

First consider the case of positive feedback (Fig. 3A). When proteins are present in the system there will be rapid switching between the *G* and *G*\* states. However, in the rare case of an extinction of proteins in the *G*\* state and where protein binding fluctuations are neglected, a transition from the bound state *G*\* to unbound state *G* does not release a protein (reaction scheme shown in Eq. (18)). The system then must wait for a protein to be produced via the low *ρ*_*u*_ firing rate if it is to leave state *G*. And hence, the waiting time for a protein to be produced at the low *ρ*_*u*_ firing rate dominates the steady-state dynamics, leading to the mode at zero (Fig. 3A red curve). However, where protein binding fluctuations are included (reaction scheme shown in Eq. (1)), a transition from the *G*\* to *G* does release a protein which can immediately bind to *G* (due to the high *σ*_*b*_ firing rate, meaning *L* ≪ 1) and hence the system does not so readily encounter an extinction of proteins (black curve Fig. 3A). We note that even for the black curve there exists a (non-visible) mode at zero, which accounts for the long waiting times in the extremely rare event (again note *σ*_*b*_ ≫ 1) that the protein released from the *G*\* state decays before binding to *G*; clearly however this is not the dominating feature where protein binding fluctuations are taken into consideration.

Now consider the case of strong negative feedback (Fig. 3B). This implies that when a protein is produced in the active *G* state (now *ρ*_*u*_ is large, *ρ*_*b*_ is very small), the rapid promoter-protein binding reaction will occur forcing the system into the *G*\* state. Where protein binding fluctuations are neglected no protein is removed upon binding and hence the number of free proteins is still 1; the system will then flip back and forth between the *G* and *G*\* states, spending (on average) more time in the *G*\* state since *σ*_*b*_ ≫ *σ*_*u*_ and hence it is unlikely more than 1 protein will ever be present (due to very small *ρ*_*b*_ and *σ*_*b*_ ≫ 1), hence the mode at *n* = 1 for the red curve in Fig. 3B. In the event of a protein extinction the unbound state *G* will quickly produce another protein in the *G* state. For strong negative feedback including the binding fluctuation, the rapid promoter-protein binding reaction will instead remove a protein from the system. Again, since it is unlikely the system will contain more than one protein (bound or otherwise), and since the system spends much more time in the *G*\* state (*σ*_*b*_ ≫ *σ*_*u*_) the probability distribution for the number of free proteins will have a mode at zero (black curve Fig. 3B).

The results stated thus far are for steady-state conditions. It would also be interesting to understand the difference between the heuristic reduced master equation Eq. (9) and the exact master equation Eq. (67) for finite time. Since this is analytically intractable we use stochastic simulations to explore this question. Figure 4 summarizes the results of such simulations for two different parameter sets: (i) 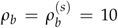 in which case the heuristic predicts a very different steady-state mean number of proteins than the exact reduced master equation; (ii) 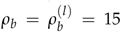 where the heuristic and exact reduced master equations are indistinguishable at steady-state (see Fig. 4(A)). In Fig. 4B we show three independent trajectories of the SSA corresponding to the the exact and heuristic reduced master equations for the two parameter choices; the vast difference between the trajectories of the heuristic and the exact for 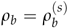 are particularly striking. In Fig. 4C we show the mean number of proteins as a function of time for the exact and heuristic reduced master equations (solid lines) and compare with the same predicted from the deterministic equations (dashed lines). Two observations can be made: (i) for both parameter sets, the deterministic reaches steady-state at a much earlier time than the stochastic models; (ii) for 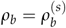 the heuristic predicts that the differences from the deterministic do not decrease with time while the exact predicts that the differences from the deterministic decrease with time (compare top two sub-figures in Fig. 4C). In contrast, for 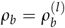 both master equations predict that differences from the deterministic decrease with time. Taken together, the results indicate that the full time-dependent solution of the heuristic reduced master equation is an accurate reflection of the exact reduced master equation provided *ρ*_*b*_, the protein production rate in state *G*\*, is large enough so that we are far away from the switching point of the positive feedback loop.

**Figure 4:**
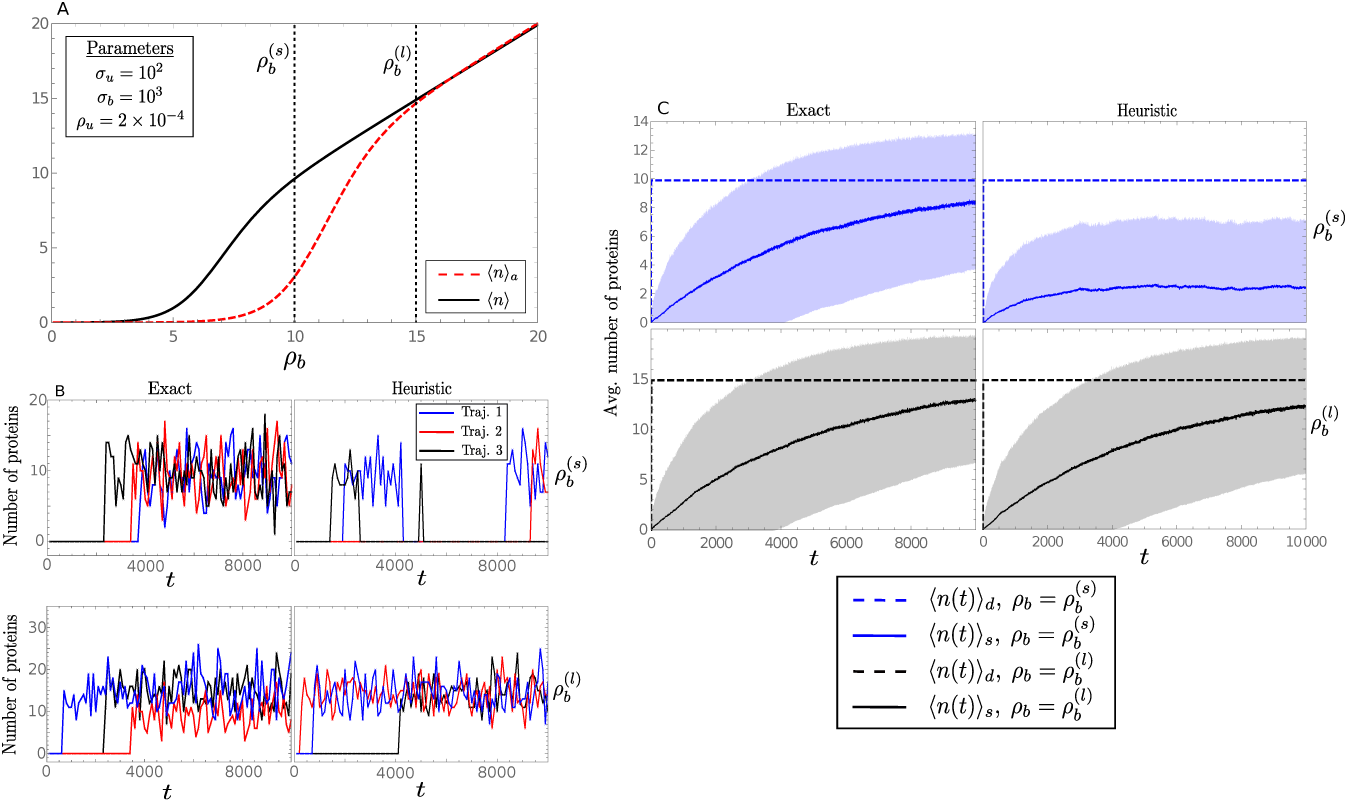
Plots comparing the time-evolution of the heuristic reduced master equation Eq. (9) and the exact master equation Eq. (67) for fast promoter switching conditions, positive feedback and small *L* (0.1). In (A) we show the switching characteristics of the positive feedback loop as a function of *ρ*_*b*_ for the two master equations in steady-state conditions. Here dotted lines define the values of 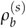 subscript *s* for small) and 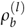 (*l* for large) used throughout the rest of the figure. In (B) we plot three independent trajectories from the SSA corresponding to the two master equations. Each trajectory shown is down-sampled 1:100 for visual clarity. The top row corresponds to 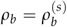 and the bottom row corresponds to 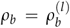. In (C) we plot the mean number of proteins as a function of time as predicted by the exact and heuristic reduced master equations (shown as solid lines and denoted by the subscript *s* in the legend) and by the deterministic rate equations (shown as dashed lines and denoted by the subscript *d* in the legend). The shaded regions show one standard deviation about the mean. The moments were calculated over 2 × 10^3^ SSA trajectories. All sub-figures compare two different parameter sets, one for small *ρ*_*b*_ (where at steady-state the heuristic and exact differ considerably) and one for large *ρ*_*b*_ (where at steady-state the differences are negligible), as indicated in Fig. 4(A). Note that min(*σ*_*u*_, *σ*_*b*_) ≫ max(1, *ρ*_*u*_, *ρ*_*b*_) and hence fast promoter switching is ensured for both parameter sets. See text for discussion.

### 2.6 Numerical computation of the distance measure between steady-state distributions

To further understand the regions of parameter space where the heuristic and exact reduced master equations differ, we numerically compute the Hellinger distance (HD) between the exact steady-state solution of the heuristic master equation Eq. (11) and the exact steady-state solution of the master equation Eq. (73) for a large region of parameter space for the positive feedback loop: *N* varying between 10^−5^ and 1, and *L* varying between 10^−4^ and 1. The results are shown as a heatmap in Fig. 5(A). Note that the Hellinger distance is a distance measure between two probability distributions; it is convenient for interpretation since the distance is a fraction, i.e. a HD value of 0 means that two distributions are identical and a HD value approaching 1 means that the distributions are very different from one another. Specifically, a maximum distance 1 is achieved when one of the distributions assigns probability zero to every set to which the other distribution assigns a positive probability. In Fig. 5(B) we calculate the Hellinger distance between the steady-state solution of the exact reduced master equation Eq. (28) and the exact steady-state solution of the master equation Eq. (73). Note that there is clear timescale separation across the whole region of parameter space used for the heatmaps as demonstrated in Fig. 5(E) and hence based on conventional wisdom, one would expect the heuristic reduced master equation to be accurate at all points in this space. However Fig. 5(A) shows this is not the case – the HD between the distributions predicted by the exact and heuristic reduced master equations varies widely between 0 and 1. In contrast Fig. 5(B) shows that the HD between the distributions predicted by the exact and exact reduced master equations is very close to zero across all of parameter space thus verifying that the latter is the correct form of the reduced master equation under fast promoter switching conditions. From Fig. 5(A) we see that there is a trend for the HD between the heuristic and exact master equations to decrease with increasing *L* which agrees with the theoretical prediction in previous sections that the differences are significant for very small *L* and disappear in the limit of large *L*. As well, there is a trend for the HD to decrease with increasing *N* and to be particularly small close to *N* = 1; this agrees with the the theoretical prediction that for *N* = 1 the heuristic and exact precisely agree because in this case there is no effective feedback mechanism.

**Figure 5:**
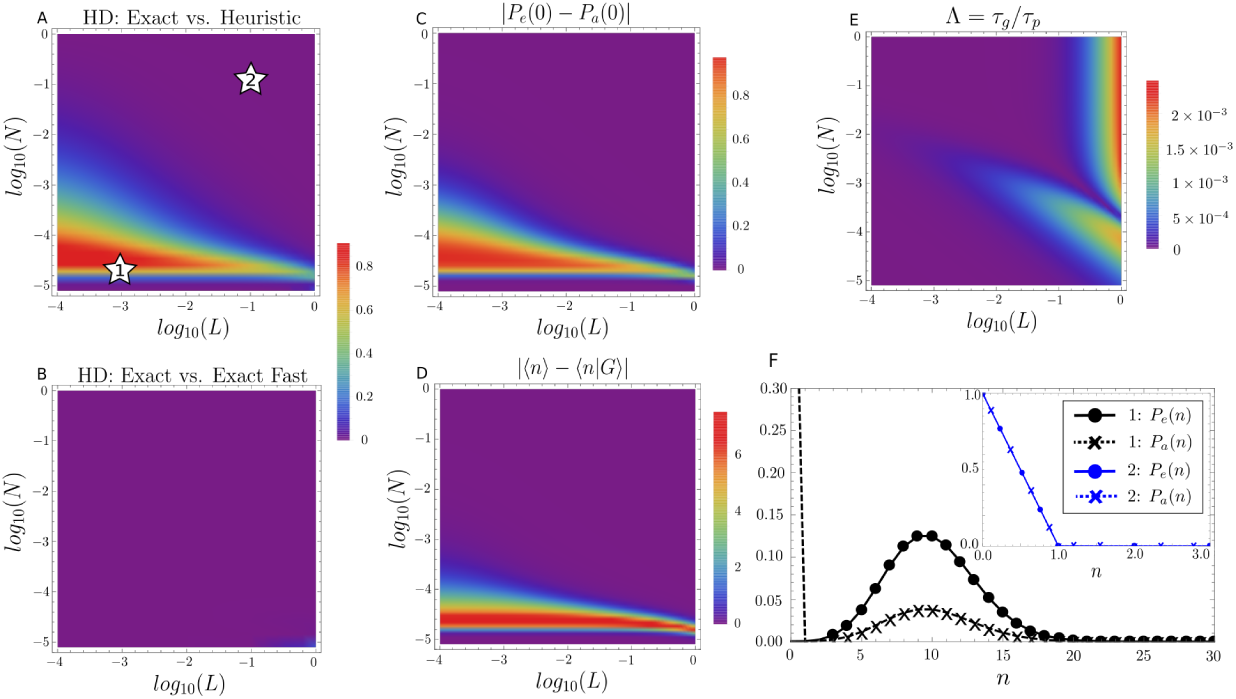
Quantifying the differences between the steady-state protein distributions predicted by the exact master equation Eq. (73), the heuristic reduced master equation Eq. (11) and the exact reduced master equation Eq. (28) for positive feedback loops. *P*_*e*_ (*n*) denotes the exact steady state solution of the exact master equation (i.e. the exact solution of the reaction scheme from Eq. (1), see derivation in Appendix B) and *P*_*a*_ (*n*) denotes the distribution from the heuristic reduced master equation. We note that *P*_*e*_ (*n*) takes the role of *P*_*FSP*_ used in previous figures. In (A) we show the Hellinger distance (HD) between the predictions of the exact master equation and heuristic reduced master equation. In (B) we show the HD between the predictions of the exact master equation and exact reduced master equation (denoted as exact fast). (C) Shows the absolute difference between the probability of zero protein molecules predicted by the exact master equation and the probability of zero protein molecules predicted by the heuristic reduced master equation. (D) Shows the absolute difference ⟨*n*⟩ and ⟨*n*|*G*⟩ computed using the exact master equation. (E) Shows that the whole region of parameter space chosen has suitable deterministic timescale separation where *τ*_*g*_ = 1/*λ*_1_, *τ*_*p*_ = 1/*λ*_2_ and *λ*_*i*_ are the eigenvalues of the Jacobian of the deterministic rate equations Eqs. (2–3) evaluated at steady-state. Note that it is expected from Fig. 4 that the stochastic timescale separation will be much greater than the deterministic. Parameters *σ*_*u*_ = 100, *ρ*_*u*_ = 2 × 10^−4^ are fixed throughout the figure, with *σ*_*b*_ varying in the range 100 − 10^6^ (small *L*) and *ρ*_*b*_ varying in the range 2 × 10^−4^ −25 (positive feedback). Numbered stars in (A) indicate the two points in parameter space whose corresponding probability distributions of protein numbers we show in (F). See text for discussion.

In Fig. 5(F) we plot the protein distributions predicted by the heuristic and exact master equations for the star points labeled (1) and (2) in Fig. 5(A) which are positioned in regions of high and low HD, respectively. Note that for point (1) the heuristic predicts that the probability that the protein numbers are zero is high whereas the the exact predicts the probability that the protein numbers are zero is very small. Inspired by this observation, as well as the theoretical prediction of modes at zero for small *L* given by Eq. (36), in Fig. 5(C) we plot a heatmap of the absolute difference between the height of the zero modes of the exact master equation and the heuristic reduced master equation and find that this heatmap is in very good agreement with the heatmap for the HD shown in Fig. 5(A). *This verifies our intuition that the differences between the protein distributions of the two master equations is mostly due to differences in their prediction of the probability of zero proteins at steady-state*. Inspired by the result in Appendix A that the deterministic time-evolution equations can be obtained from the full master equation under the assumption ⟨*n*|*G*⟩ ≈ ⟨*n*⟩ (which is equivalent to independence of protein and gene fluctuations), we plot in Fig. 5(D) a heatmap of | ⟨*n*| *G*⟩ − ⟨*n*⟩| which also shows broad similarity to that in Fig. 5(A).

In Fig. 6 we show the results of the same analysis as in Fig. 5 but now for the case of negative feedback loops. As before, the largest HD between the heuristic reduced master equation and the exact master equation is found for *N* far away from the trivial case of *N* = 1 and for small *L*, in line with the theoretical predictions of the previous section. Both the heatmaps of the absolute difference between the heights of the zero modes (Fig. 6(C)) and of the absolute difference between the protein mean and the conditional protein mean (Fig. 6(D)) show high correlation with the HD heatmap in Fig. 6(A). The differences between the heuristic and exact protein distributions for the large HD and small HD in Fig. 6(A) (star point 1, 2 respectively) are shown in Fig. 6(F). Note that the differences between the two cases amount to the order of a single molecule and are far smaller than the differences found for positive feedback (compare Fig. 5(F)), verifying the theoretical predictions of the previous section, namely that the heuristic fails worst for positive feedback.

**Figure 6:**
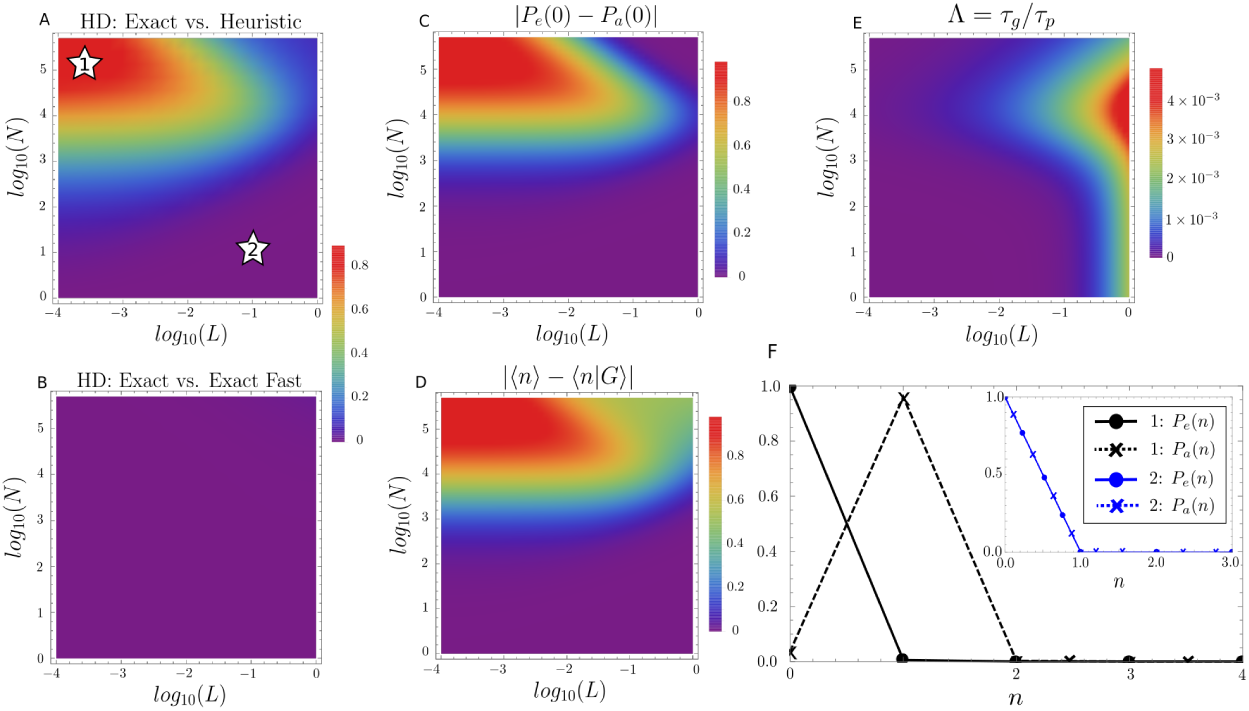
Quantifying the differences between the steady-state protein distributions predicted by the exact master equation Eq. (73), the heuristic reduced master equation Eq. (11) and the exact reduced master equation Eq. (28) for negative feedback loops. In (A) we show the Hellinger distance (HD) between the predictions of the exact master equation and heuristic reduced master equation. In (B) we show the HD between the predictions of the exact master equation and exact reduced master equation. (C) Shows the absolute difference between the probability of zero protein molecules predicted by the exact master equation and the probability of zero protein molecules predicted by the heuristic reduced master equation. (D) Shows the absolute difference ⟨*n*⟩ and ⟨*n*|*G*⟩ computed using the exact master equation. (E) Shows that the whole region of parameter space chosen has suitable deterministic timescale separation where *τ*_*g*_ = 1/*λ*_1_, *τ*_*p*_ = 1/*λ*_2_ and *λ*_*i*_ are the eigenvalues of the Jacobian of the deterministic rate equations Eqs. 2–3) evaluated at steady-state. Note that it is expected from Fig. 4 that the stochastic timescale separation will be much greater than the deterministic. Parameters *σ*_*u*_ = 100 and *ρ*_*b*_ = 2 × 10^−4^ are fixed throughout the figure, with *σ*_*b*_ varying between 100 and 10^6^ (small *L*) and *ρ*_*u*_ varying between 2 × 10^−4^ and 25 (negative feedback). Numbered stars in (A) indicate the two points in parameter space whose corresponding probability distributions of protein numbers we show in (F). See text for discussion.

### 2.7 Extending results to the case of multiple protein binding

Here we briefly treat the more general case where multiple protein molecules can bind the promoter, a common case in nature often associated with cooperative behaviour. The reaction scheme is an extension of (1) and reads:

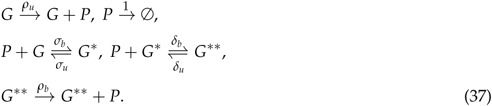

Writing the deterministic rate equations for this system and making the assumption of fast promoter switching such that *∂*_*t*_ ⟨*g*⟩_*d*_ ≈ 0, *∂*_*t*_ ⟨*g*\* ⟩_*d*_ ≈ 0 and *∂*_*t*_ ⟨*g*\*\*⟩_*d*_ ≈ 0 it is straightforward to show that the effective deterministic time-evolution equation for the protein numbers has the form:

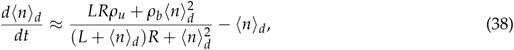

where *L* = *σ*_*u*_ /*σ*_*b*_ and *R* = *δ*_*u*_ /*δ*_*b*_. It follows by the same reasoning as in Section 2.2 that the corresponding heuristic reduced master equation for protein dynamics is given by Eq. (9) with the effective propensities:

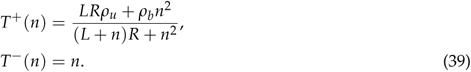

In the limit of small *L* and *R*, Eq. (39) reduces to the effective propensities given by Eq. (14) while in the limit of large *L* and *R*, Eq. (39) reduces to the effective propensities given by Eq. (12). Hence the solutions of the heurestic master equation in these two limits are given by Eq. (17) and Eq. (13).

By inspection of the reaction scheme (37) it is obvious that for small *L* and *R*, the gene will be mostly in state *G*\*\* and hence the principal reactions determining the protein dynamics are 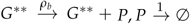. By the same reasoning, it follows that for large *L* and *R*, the gene will be mostly in state *G* and the principal reactions are 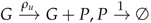. Hence the solution of the exact master equation of in the limit of small and large *L, R* is Poisson and given by Eq. (32) and Eq. (31) respectively. *Hence all the conclusions previously reached regarding the differences between the heuristic reduced master equation and the exact master equation for single protein binding for the case of small and large L also hold for multiple protein binding for the cases of small and large L, R*. Note that the derivations here assume the exchangeability of the limits of large/small *L, R* and large time; hence the proof here presented is not formal but the results are the expected ones and are confirmed by simulations (see later).

It is interesting to find the general conditions under which the heuristic reduced master equations generally agrees with the exact. For the case of single promoter binding, we showed in Appendix A that the deterministic rate equations agreed with the mean of the exact master equation when ⟨*n*⟩ = ⟨*n* |*G*⟩. Next we derive a similar condition for the case of multiple protein binding. We start by noting that under fast promoter switching conditions, the reactions 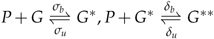 are in equilibrium and hence the deterministic rate equations yield:

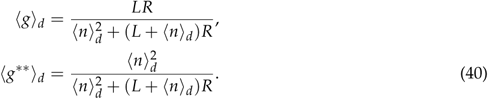

Next we derive equations for the same quantities but from the exact master equation (denoted ⟨*g*⟩, ⟨*g*\*⟩ and ⟨*g*\*\*⟩). Writing the master equation for the same two reversible reactions in equilibrium, one can deduce the moment equations:

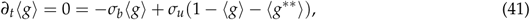

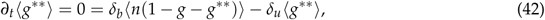

from which we can deduce:

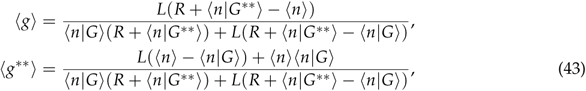

where we used the definitions of conditional means: ⟨*n*|*G*⟩ = ⟨*ng*⟩/⟨*g*⟩ and ⟨*n*|*G*\*\* ⟩ = ⟨*ng*\*\* ⟩ / ⟨*g*\*\* ⟩. Comparing Eq. (40) and Eq. (43), we see that they can only be equal if the following condition is true:

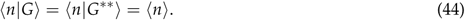

By means of the definition of the mean in terms of conditional means ⟨*n*⟩ = ⟨*n*|*G*⟩⟨*g*⟩ + ⟨*n*|*G*\* ⟩⟨*g*\*⟩+ +⟨*n*|*G*\*\*⟩ ⟨*g*\*\*⟩, one can deduce the final condition required for the agreement of the time-evolution equations for the mean protein number according to the deterministic rate equations and the exact master equation:

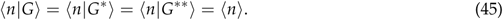

Since the heuristic is based on the deterministic, we expect that this condition is also an indicator of when the heuristic and exact master equations agree. In Fig. 7 we verify this intuition using simulations: when the above condition is approximately met then the heuristic and exact master equations predict very similar distributions of protein numbers (see Fig. 7(A),(C)) whereas the largest differences between the two master equations (see Fig. 7(B),(D)) correlate with significant differences between the three conditional mean protein numbers ⟨*n*|*G*⟩, ⟨*n*|*G*\*⟩, ⟨*n*|*G*\*\*⟩.

**Figure 7:**
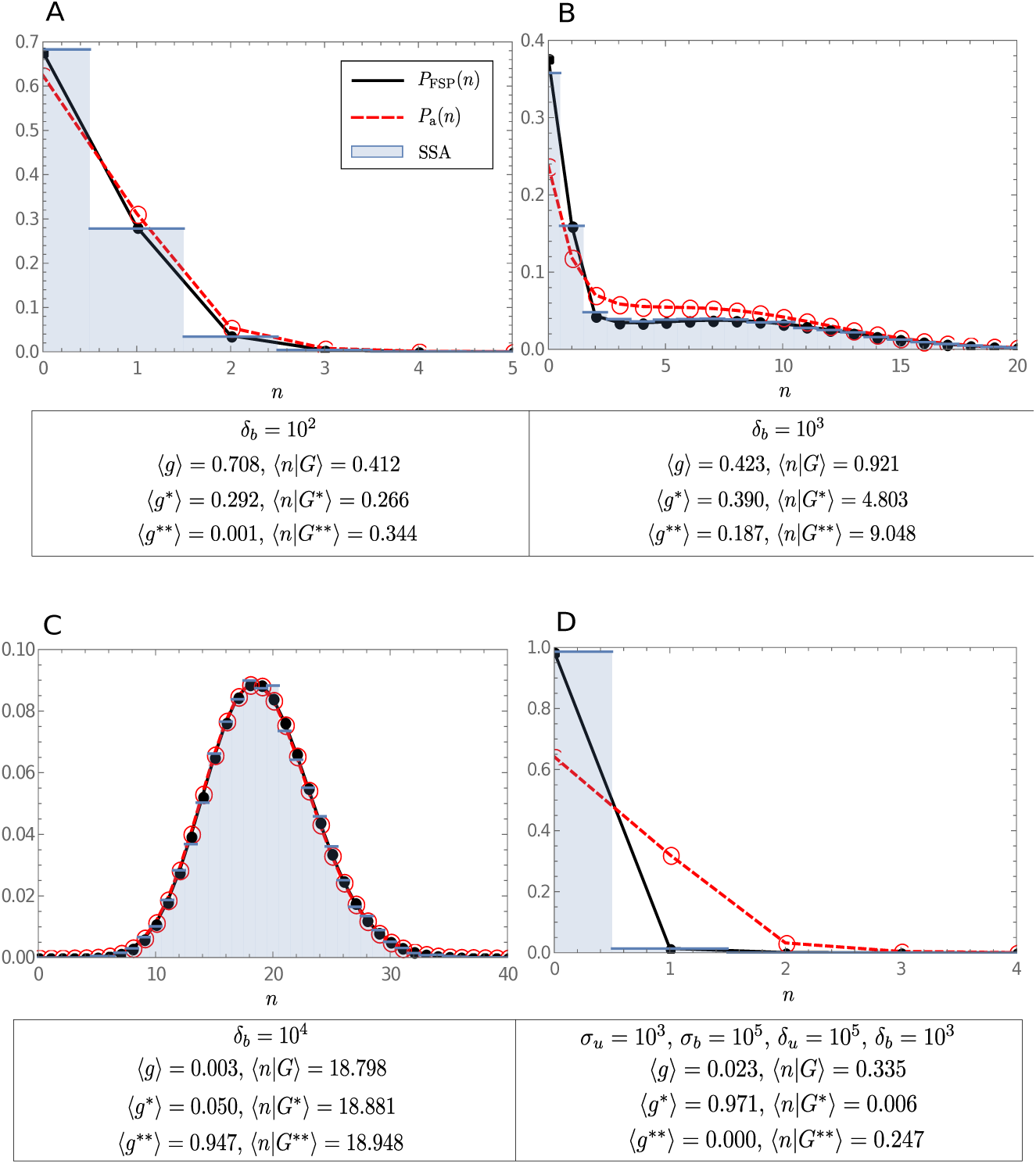
Plots comparing the heuristic master equation (Eq. (9) with Eq. (39)) and exact master equation predictions for the protein number distributions of a positive feedback loop (*ρ*_*u*_ = 0.5, *ρ*_*b*_ = 20) with multiple protein binding (37). All calculations done using FSP. Fast promoter switching is enforced by choosing min(*σ*_*u*_, *σ*_*b*_, *δ*_*u*_, *δ*_*b*_) ≫ max(1, *ρ*_*u*_, *ρ*_*b*_). In (A)-(C) parameter values are fixed to *σ*_*u*_ = 10^4^, *σ*_*b*_ = 10^4^, *δ*_*u*_ = 10^4^ and *δ*_*b*_ is varied in the range 10^2^ − 10^4^. Plot (A) shows a case where the system spends most of its time in state *G*, in (B) we show a case for which all states *G, G*\* and *G*\*\* are frequently accessed by the system and in (C) we show a case where the state *G*\*\* dominates. Sub-figure (D) shows a case where the systems spends most of its time in state *G*\*. Note that the differences between the heurestic and exact master equation are reflected in the differences between the values of the mean number of proteins conditional on each state, with the smallest differences occurring for cases (A) and (C), in agreement with the condition given by Eq. (45).

## 3 Model reduction for bursty feedback loops

In this section we consider model reduction for feedback loops in which there is an implicit mRNA description. It has been rigorously shown that when mRNA degrades much quicker than proteins (a common situation for bacteria and yeast cells) then the mRNA does not need to be explicitly described but rather implicitly manifests through protein bursts [50]. Studies have elucidated the implications of taking into account protein bursting on downstream pathways and shown its importance [51]. Hence we now consider a feedback loop with an implicit mRNA description which has the effective reaction scheme:

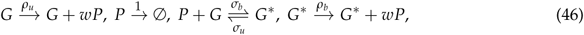

where *w* is the protein burst size which is a random positive integer drawn from the geometric distribution with mean *b* = *k*/*d*_*M*_, *k* is the rate at which mRNA is translated into proteins and *d*_*M*_ is the mRNA degradation rate. Note that the geometric form of the protein burst distribution has been shown theoretically [52] and verified experimentally [53].

### 3.1 Heuristic stochastic model reduction

We are interested in a reduced description of protein fluctuations in the limit of fast promoter switching. Clearly the effective master equation has to have the general form:

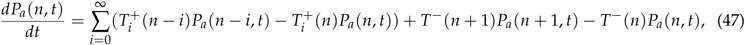

where 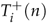 is the probability, given *n* proteins, that a protein burst of size *i* will occur in the time interval [*t, t* + *dt*) and *T*^−^ (*n*)*dt* is the probability, given *n* proteins, that a protein degradation event reducing the number of proteins by one will occur in the time interval [*t, t* + *dt*). Next we use the deterministic rate equations to guess the equations for 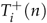 and *T*^−^ (*n*). The deterministic rate equations corresponding to (46) are given by:

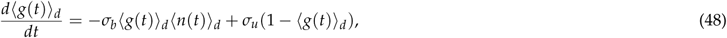

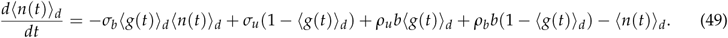

Note that these equations are the same as the deterministic rate equations for the non-bursty case given by Eqs. (2–3) except that *ρ*_*u*_ is replaced by *ρ*_*u*_*b* and *ρ*_*b*_ is replaced by *ρ*_*b*_*b*; this directly follows from the definition of *b* as the mean protein burst size. Assuming fast promoter switching, *∂*_*t*_ ⟨*g*(*t*)⟩_*d*_ ≈ 0, it follows that an effective reduced rate equation for the mean protein numbers is:

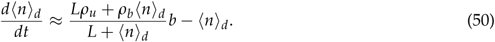

The form of this effective equation combined with the fact that we know that burst size is distributed according to a geometric distribution with mean *b* suggests a one-variable master equation of the form Eq. (47) with effective propensities:

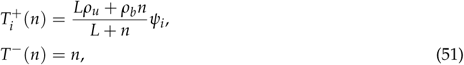

where *ψ*_*i*_ is the probability that a burst has size *i* which is given by *b*^*i*^ /(1 + *b*)^*i*+1^. If we denote the angled brackets with subscript *a* as the statistical averages calculated using the heuristic master equation Eq. (47) with propensities given by Eq. (51) then it follows that:

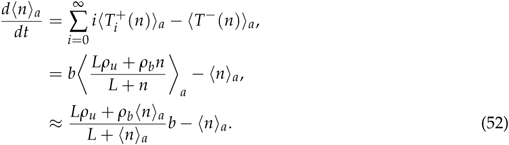

Note that this equation is the same as the reduced rate equation Eq. (50) (upon replacing ⟨*n*⟩ _*a*_ by ⟨*n*⟩) and hence verifies that the form of the effective propensities given by Eq. (51) guarantees equivalence between the effective equation for the time-evolution of the mean protein numbers of the heuristic master equation and the reduced deterministic rate equation in the limit of small protein number fluctuations when ⟨*n*⟩ _*a*_ ≈ *n*.

The heuristic stochastic model given by Eqs. (47,51) is difficult to solve exactly in steady-state because there are no known general solutions for one species reaction systems with multi-step reactions, i.e. reactions leading to the production of more than one molecule at a time [44]. However, as we now show, provided we can assume exchangeability of the limits of large/small *L* and large time then closed-form solutions can be obtained for the steady-state distributions of protein numbers.

#### 3.1.1 The limit of large *L*

In this limit, Eq. (51) reduces to the simpler form:

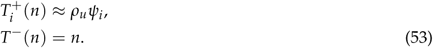

Substituting these in the heuristic reduced master equation Eq. (47), multiplying both sides by *z*^*n*^ and taking the sum over *n* on both sides of this equation we get the generating function equation:

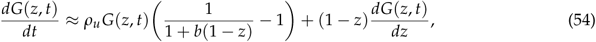

where *G*(*z, t*) = Σ_*n*_ *z*^*n*^ *P*_*a*_ (*n, t*). This equation can be solved in steady-state yielding *G*(*z*) = (1 − *b*(*z* − 1))^−*ρu*^ which implies that:

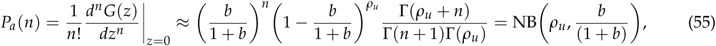

where NB(*x, y*) stands for a negative binomial distribution with parameters *x, y* and mean *xy*/(1 − *y*).

#### 3.1.2 The limit of small *L*

In this limit, Eq. (51) reduces to the simpler form:

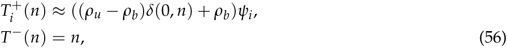

where *δ*(0, *n*) is the Kronecker delta. Substituting these in the heuristic reduced master equation Eq. (47), multiplying both sides by *z*^*n*^ and taking the sum over *n* on both sides of this equation we get the corresponding generating function equation:

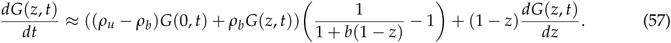

In steady-state, this equation has the solution:

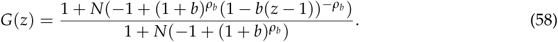

Hence the steady-state probability distribution is given by:

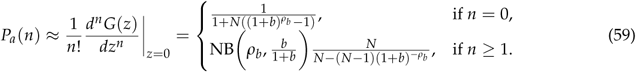

### 3.2 Exact stochastic model reduction

To determine how accurate is the heuristic model reduction we need to compare it with the reduction done on the exact model in the limit of fast promoter switching. Unlike the case of a non-bursty feedback loop, the exact solution of the the chemical master equation for reaction scheme (46) is unknown. However by taking the same approach as we did in Section 2, it is easy to find the the solution of the chemical master equation for the case of fast promoter switching and *L* being either very small or very large.

The limit of fast promoter switching implies that the reaction 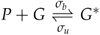 in the reaction scheme (46) is approximately in equilibrium for all times. From the chemical master equation for this reversible reaction one finds that the fraction of time that the gene is ON is given by Eq. (71). Hence it follows that in the limit of small *L*, ⟨*g*⟩ is also very small, the gene spends most of its time in state *G*\* and consequently the only effective reactions determining the protein dynamics are:

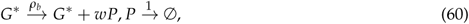

where *w* is the protein burst size which is a random positive integer drawn from the geometric distribution with mean *b*. The chemical master equation for these two reactions can be easily solved in steady-state leading to a negative binomial distribution, *P*(*n*) ≈ NB(*ρ*_*b*_, *b*/(1 + *b*)). In the opposite limit of large *L*, ⟨*g*⟩ is approximately 1, the gene spends most of its time in state *G* and consequently the only effective reactions determining the protein dynamics are:

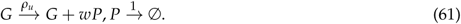

Solving the chemical master equation in steady-state (using the generating function method) for these two reactions leads to another negative binomial solution, *P*(*n*) ≈ NB(*ρ*_*u*_, *b*/(1 + *b*)).

Hence summarizing the results of Sections 3.1 and 3.2, we can state that the heuristic and exact stochastic model reduction in the limit of fast promoter switching agree for large *L* (both predict a negative binomial distribution, *P*(*n*) = *P*_*a*_ (*n*) = NB(*ρ*_*u*_, *b*/(1 + *b*))) but disagree for small *L*: the exact reduction predicts a negative binomial distribution, *P*(*n*) = NB(*ρ*_*b*_, *b*/(1 + *b*)) while the heuristic reduction predicts the different distribution given by Eq. (59). These results qualitatively parallel those previously obtained for a non-bursty feedback loop.

To further understand the differences between these two distributions for small *L* we now look at the mean protein numbers, the Fano Factor and the Coefficient of Variation of protein number fluctuations:

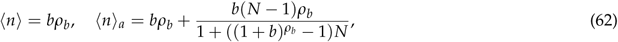

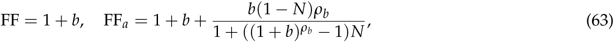

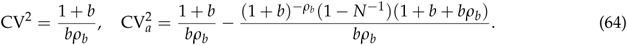

Hence for *N* < 1 (positive feedback), the heuristic underestimates the mean protein number and over-estimates the Fano Factor and the Coefficient of Variation of protein number fluctuations and the opposite occurs when *N >* 1 (negative feedback). The deterministic rate equations predict a mean of *bρ*_*b*_ (can be deduced from Eq. (50) in limit of small *L*) which agrees with ⟨*n*⟩ but not with ⟨*n* ⟩_*a*_ and hence the heuristic artificially predicts noise-induced deviations from the deterministic mean. These are the same conclusions that we reached in Section 2.4 for the case of a non-bursty feedback loop. The exact reduction predicts super-Poissonian fluctuations (FF *>* 1) while the heuristic predicts the same for *N* < 1 and either super- or sub-Poissonian fluctuations for *N >* 1 (FF_*a*_ *>* 1 and FF_*a*_ < 1 respectively). We note that the prediction of sub-Poissonian fluctuations is a surprising illogical output of the heuristic model since naturally the production of proteins in bursts has to lead to number distributions which are wider than Poisson. The theoretical predictions for the mean, FF, CV are corroborated using FSP in Fig. 8(A)-(C).

**Figure 8:**
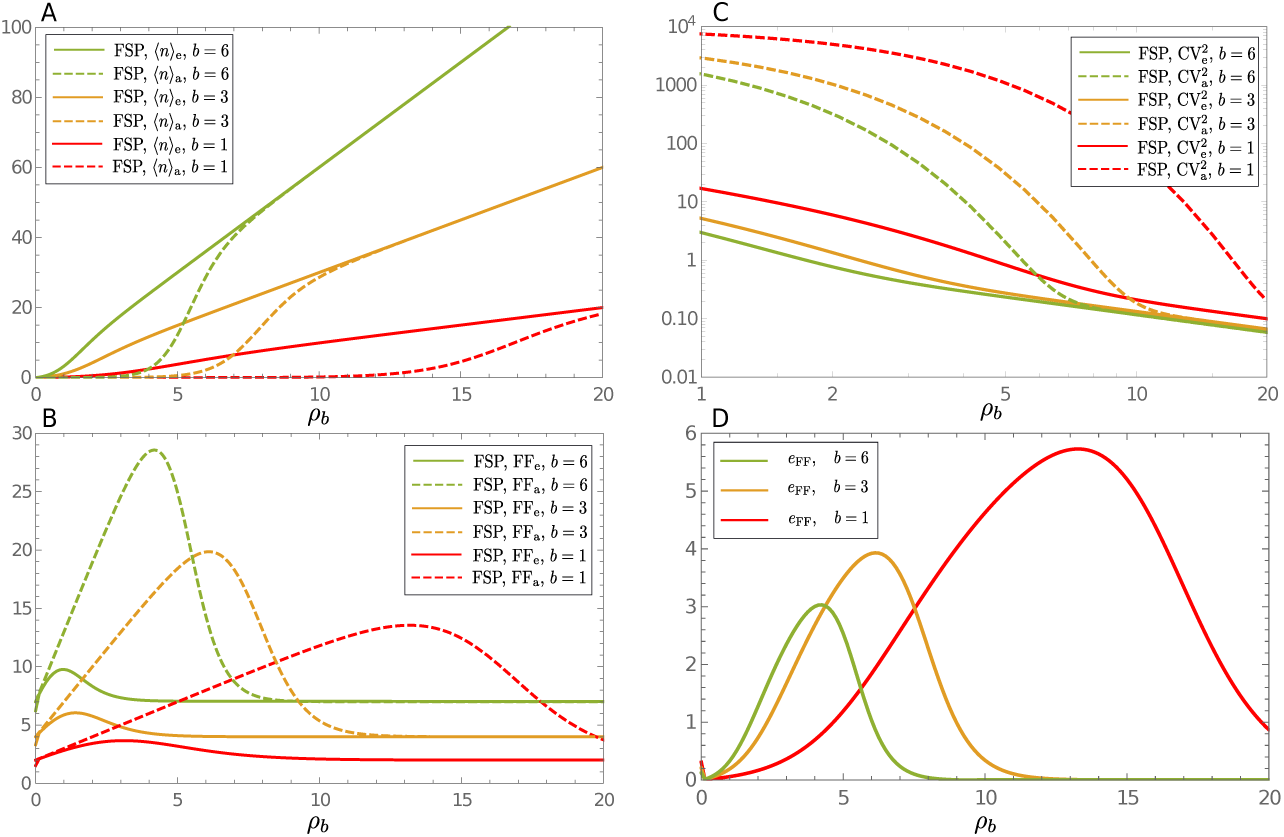
Plots showing the breakdown of the heuristic reduced master equation for fast promoter switching in the limit of small *L* for a bursty positive feedback loop. The plots show the mean protein number number (A), the Fano Factor of protein number fluctuations (B) and the Coefficient of Variation squared (C) as a function of *ρ*_*b*_ and the mean burst size *b*. In these plots we compare the FSP solution of the full master equation corresponding to reaction scheme (46) with the FSP solution of the heuristic reduced master equation Eq. (47) with Eq. (51); the latter is distinguished from the former by the subscript e. These are in good agreement with the moments calculated in the limit of small *L* and given by Eq. (62)-(64) – for example the means of the exact solution in (A) are very well approximated by *⟨n* ⟩= *bρ*_*b*_. In (D) we show the relative error in the heuristic reduced master equation’s FF computed using the data in (B) where *e*_FF_ = |FF_*e*_ − FF_*a*_| /FF_*e*_. Note that the relative error decreases with increasing mean burst size *b*. In all cases *ρ*_*u*_ = 0.0002, *σ*_*u*_ = 100 and *σ*_*b*_ = 10^5^ were chosen such that *L* is small, there is positive feedback and fast promoter switching is guaranteed.

The relative errors (made by the heuristic reduction method) for the mean, FF and CV^2^ can be computed using *e*_*m*_ =| ⟨*n*⟩ −⟨*n*⟩ _*a*_ / ⟨*n*⟩, *e*_FF_ = |FF FF_*a*_| /FF and 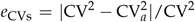, respectively. The errors for the bursty feedback loop (computed using Eqs. 62–64)) are smaller than the errors for the non-bursty feedback loop (computed using Eqs. 33–35)) provided the mean burst size *b* ≫ 1. *Hence a major prediction of our theory is that bursts in protein expression generally reduce the size of the discrepancies between heuristic and exact stochastic model reduction in the limit of small L*. This theoretical prediction is verified using FSP in Fig. 8(D).

Finally we compute the conditions for the existence of a mode of the probability distribution at *n* = 0 using the distribution obtained from the exact method *P*(*n*) = NB(*ρ*_*b*_, *b*/(1 + *b*)) and the distribution from the heuristic reduction Eq. (59) respectively:

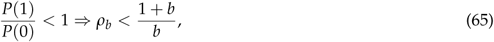

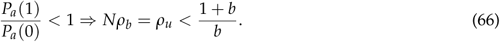

This implies that if *ρ*_*u*_ < (1 + *b*)/*b, ρ*_*b*_ *>* (1 + *b*)/*b* (a special case of positive feedback), the approximate heuristic master equation predicts an artificial mode at *n* = 0 whereas if *ρ*_*u*_ *>* (1 + *b*)/*b, ρ*_*b*_ < (1 + *b*)/*b* (a special case of negative feedback), the approximate heuristic master equation misses to predict an actual mode at *n* = 0.

## 4 Conclusion

In this paper we have conclusively shown that heuristic stochastic models with Hill-type propensities for transcriptional regulation are not generally valid under fast promoter switching conditions, as commonly assumed. Rather we show that they are valid only over a subset of parameter space consistent with the fast promoter switching assumption, namely when the rate of protein-DNA binding reaction is much less than the unbinding reaction. Our work shows that when this condition is not met, the protein distributions predicted by the heuristic models can be considerably different than the true protein distributions. These differences exist for both negative and positive feedback loops but are particularly pronounced for the latter – in this case we have shown that the heuristic model can predict an artificial mode at zero proteins, an incorrect switching point from low to high protein expression as a parameter is varied, artificial deviations of the mean number of proteins from that predicted by the rate equations and a huge overestimation of the size of protein number fluctuations and of the Fano Factor. Surprisingly, we found that the heuristic solution exactly corresponds to the fast gene switching limit of the autoregulatory system that ignores protein number fluctuations due to the protein-promoter binding reaction. Our work further builds on previous work by other authors [40, 41, 54] but has the advantage of using theory to precisely deduce the region of validity of the heuristic approach.

A number of open answered questions remain: (i) Is there a simple way of constructing a different type of reduced stochastic model which avoids the pitfalls of the common heuristic models and which also avoids the use of sophisticated mathematical analysis to derive it? The requirement of simplicity is essential because typically only such methods are widely adopted and indeed this is a main reason why the problematic heuristic reduced stochastic models treated in this paper are so widespread. (ii) What would be the differences between heuristic and correctly reduced stochastic spatial models of genetic feedback loops in the limit of fast gene switching? Would the differences between the two models increase or decrease with the diffusion coefficient of protein molecules? Spatial modeling of such systems is relatively rare but recent work in this direction [55–59] shows that these models are richer in complex behavior than their non-spatial counterparts and of course they are closer to reality. (iii) Say one used a heuristic reduced stochastic model to construct a likelihood function and then use the latter within a Bayesian approach to infer parameters of auto-transcriptional feedback loops from experimental data: how would these differ from parameters inferred using a likelihood built from a non-reduced model? A recent study [60] shows that inference from moment-based approaches is very sensitive to the type of approximation used to construct the likelihood function and hence suggests large differences between the parameters inferred using heuristic reduced or exact master equations. In conclusion, our study shows that care must be exerted in the interpretation of the results of heuristic stochastic models.

## Author Contributions

J. H. implemented and ran all simulations, produced the figures and performed some of the calculations. R. G. conceived the study, directed it, performed most of the calculations and wrote the paper. All the authors reviewed and edited the final manuscript.

## Acknowledgments

This work was supported by a BBSRC EASTBIO PhD studentship to J. H and BBSRC grant BB/M025551/1 to R. G.

## A Relationship between the deterministic and stochastic models

For the non-bursty feedback loop (1), the stochastic description is given by the chemical master equation which can be formulated as a set of two coupled equations:

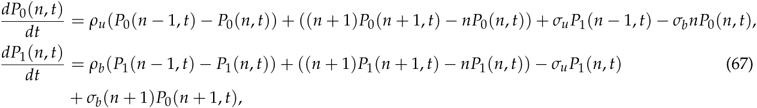

where *P*_0_(*n, t*) is the probability that at time *t* there are *n* proteins and the gene is in state *G* while *P*_1_(*n, t*) is the probability that at time *t* there are *n* proteins and the gene is in state *G*\*. Note that time *t* is non-dimensional and equal to the actual time multiplied by the protein degradation rate. The probability of *n* proteins is then given by *P*(*n, t*) = *P*_0_(*n, t*) + *P*_1_(*n, t*). Using these equations it is straightforward to show that the time-evolution equations for the first moments:

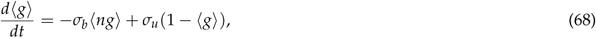

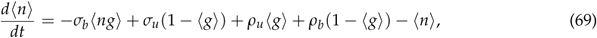

where ⟨*n*⟩ = ∑_*n*_ *nP*(*n*) is the mean numbers of proteins, ⟨*g*⟩ = ∑_*n*_ *P*_0_(*n*) is the fraction of time the gene is in the ON state (or equivalently the average number of gene in ON state) and⟨*ng*⟩ = ∑_*n*_ *nP*_0_(*n*). Note that we have here suppressed the time dependence for convenience. A comparison of Eqs. (2)-(3) with Eqs. (68)-(69) shows that the two are the same if ⟨*ng*⟩ = ⟨*n*⟩ ⟨*g*⟩, i.e. the deterministic and exact stochastic models agree in the means if the fluctuations in the gene and protein numbers are independent of each other.

We next show that in the limit of fast promoter switching and when there is a non-zero correlation between the fluctuations of protein and gene, the exact stochastic model gives a time-evolution equation for the protein number mean which is different than that given by the deterministic analysis. The limit of fast promoter switching implies that *d* ⟨*g*⟩ /*dt* ≈ 0 and hence using Eq. (68) it follows that the mean number of proteins conditional on the gene being in state *G* can be written as:

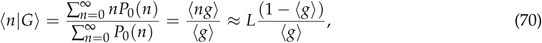

from which we obtain after rearrangement:

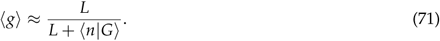

Substituting in Eq. (69) we obtain an effective equation for the time-evolution of the protein numbers under the condition of fast promoter switching:

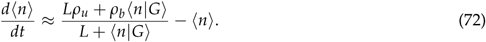

Contrasting this equation with the effective equation obtained through the deterministic approach, Eq. (4) we see that the two are generally different. They are only the same when ⟨*n|G*⟩ = ⟨*n*⟩, i.e. the mean number of proteins conditional on the gene being in state *G* is equal to the mean number of proteins which occurs when gene and protein number fluctuations are independent.

## B Exact steady-state solution of non-bursty feedback loop in limit of fast gene switching

The exact steady-state solution of Eq. (67) has been previously reported in the literature [7, 61] and is given by:

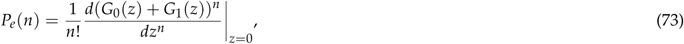

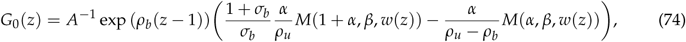

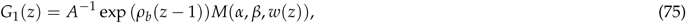

with the definitions:

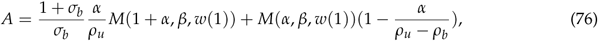

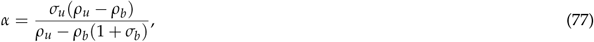

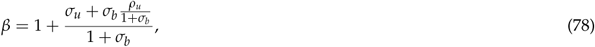

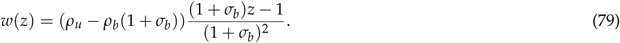

Note that *M*(*x, y, z*) is the Kummer confluent hypergeometric function and *G*_*i*_ (*z*) is the generating function ∑_*n*_ *z*^*n*^ *P*_*i*_ (*n*). Replacing *σ*_*u*_ by *σ*_*u*_ /*ϵ* and *σ*_*b*_ by *σ*_*b*_ /*ϵ* and taking the limit of *ϵ* → 0 (the fast switching limit), we find that *G*(*z*) becomes a function of only three non-dimensional parameters *L* = *σ*_*u*_ /*σ*_*b*_, *N* = *ρ*_*u*_ /*ρ*_*b*_, *ρ*_*b*_ and the corresponding steady-state distribution of protein numbers (to leading-order in *ϵ*) has the form:

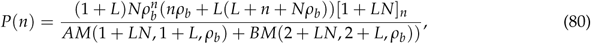

where

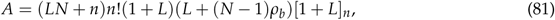

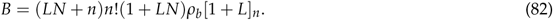

## C Limits of small and large *L* from exact steady-state solutions

### C.1 Interchanging the sum and the limit

Here we prove a result which will be used in Sections C.2 and C.3, for the purpose of interchanging limits of *L* with the infinite sum that defines the Kummer function (for example that in Eq. (11) and Eq. (28)). For reference, the Kummer function is defined through the sum:

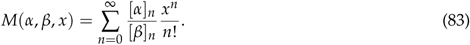

We will prove that:

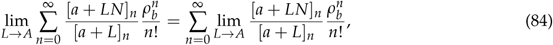

where *A* is some real number.

Let *f*_*n*_ = [*a* + *LN*]_*n*_ /[*a* + *L*]_*n*_ which implies *f*_*n+*1_ = *f*_*n*_ (*a* + *n* + *LN*)/(*a* + *n* + *L*). Consider first the case *N ≥* 1. Since (*a* + *n* + *LN*)/(*a* + *n* + *L*) ≤ *N* then if *f*_*n*_ ≤ *N*^*n*^ this implies that *f*_*n+*1_ ≤ *N*^*n*+1^. Also it is easy to check that *f*_1_ ≤ *N*. Hence by induction it follows that if *N ≥* 1 then *f*_*n*_ ≤ *N*^*n*^. Similarly it is straightforward to prove by induction that if *N* < 1 then *f*_*n*_ < 1. It then follows that:

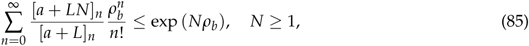

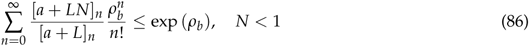

Since the sums are bounded by a finite value, it follows by the dominated convergence theorem that the limit and sum can be switched, i.e. Eq. (84) holds.

### C.2 The limit of large *L*

Making use of a standard result [62]:

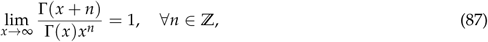

and [*x*]_*n*_ = Γ[*x* + *n*]/Γ[*x*], we have that for any *a* ∈ ℝ:

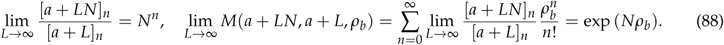

Note that here we have used the interchange of sum and limit as given by Eq. (84). Using Eq. (88) and *N* = *ρ*_*u*_ /*ρ*_*b*_ it follows that in the limit of large *L*, the steady-state distribution of molecule numbers as predicted by the reduced master equation Eq. (11) and by the full master equation in the limit of fast promoter switching Eq. (28) is a Poissonian with mean *ρ*_*u*_:

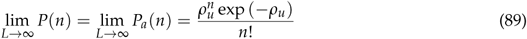

### C.3 The limit of small *L*

Using the definition of the Pochhammer symbol and the standard result Γ(*x*) ~ 1/*x* as *x* → 0, one can show that:

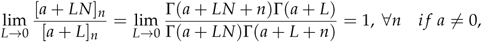

and

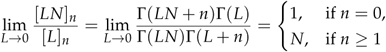

Using these two results, it then follows that:

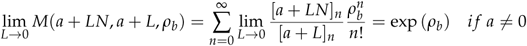

and

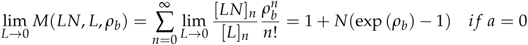

Note that here we have used the interchange of sum and limit as given by Eq. (84). Using these results it is straightforward to show that in the limit of small *L*, the steady-state distribution of molecule numbers predicted by the heuristic master equation Eq. (11) reduces to:

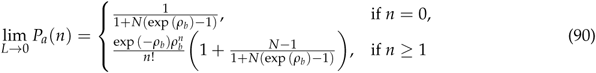

while the steady-state distribution of molecule numbers predicted by the master equation in the limit of fast promoter switching Eq. (28) reduces to:

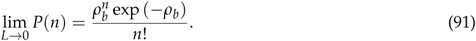

Clearly *P*(*n*) ≠ *P*_*a*_ (*n*) since the latter is not-Poissonian and hence shows that in the small *L* limit, the approximate heuristic master equation gives the incorrect answer.

